# Molecular regulation of *Bna1205ams1* required for male fertility and development of a recessive genic male-sterility system in *Brassica napus*

**DOI:** 10.1101/2024.02.25.581914

**Authors:** Lijing Xiao, Jinze Zhang, Kunjiang Yu, Xu yang, Qian Wang, Hairun Jin, Qingjing Ouyang, Entang Tian

## Abstract

Through the comprehensive use of two pollination control systems of cytoplasmic male sterility (CMS) and genic male sterility (GMS), the rapeseed yield was increased by more than 20%. However, more hybrid production systems and detailed mechanisms underlying male sterility are required. Here, we reported a novel two-line hybrid production system of 1205A for GMS and also investigated the underlying mechanism for male sterility. Five co-segregated kompetitive allele specific PCR (KASP) markers were developed and validated, which could be used for transferring the male sterility trait into external *B. napus* breeding lines and developing further two-line hybrid production systems of GMS. Inheritance studies detected one gene locus of *Bna1205ams1* for regulating the male sterility of 1205A. As a potential candidate gene of *Bna1205ams1*, *BnaC03g27700D* was fine mapped and narrowed down to a 181.47 kb region on chrC03 and validated by functional analysis. The mutation of *BnaC03g27700D* in 1205A resulted in large metabolic fluctuations, most of which were involved in aborted tapetal PCD, which could lead to reduced pollen fertility with abnormal pollen exine. The developed new GMS line of 1205AB provided us with the opportunity to identify a new male sterility gene of *BnaC03g27700D* in *B. napus*. The study of *BnaC03g27700D* aims to renew the annotation of the gene and provide new resources for basic research on the genetic control of male sterility.

## Introduction

Rapeseed (*Brassica napus* L., AACC, *2n*=38) is today one of the most important oil seeds for vegetable oil production worldwide. The widespread use of hybrid breeding strategies has increased the yield of *B. napus* by more than 20% (Gupta, 2007). In China, hybrid seeds account for more than 75% of the total cultivated area (Fu, 2009). Currently, two pollination control systems, cytoplasmic male sterility (CMS) and genic male sterility (GMS), are used in *B. napus* for the development of hybrid varieties. CMS systems are still the most widely used breeding system in hybrid production, such as the Ogura system (Feng et al., 2009; Hu et al., 2008; Prakash et al., 1998) and the Polima system (Fu, 2009). Another system of genic male sterility (GMS) has attracted the enthusiasm of many rapeseed breeders due to its pure and stable male sterility, which is not affected by the cytoplasm, and its simplified breeding system from three-line to two-line, compared to CMS system. In particular, RGMS lines have been found in numerous species, such as: *B. napus* (FX et al., 1993; Li et al., 2012b; Yi et al., 2006), *Brassica oleracea* (Guo et al., 2016; Ji et al., 2017), *Gossypium hirsutum* (Raja et al., 2018; Wang et al., 2016), *Cryptomeria japonica* (Mishima et al., 2018), *Cucumis sativus* (Han et al., 2018), and *Capsicum annuum* (Cheng et al., 2018). In *B. napus*. Many different RGMS systems have been found and studied in detail, which are controlled by different genes. A spontaneous mutant was found from a distant hybridization between *B. napus* and *B. rapa*, and one gene locus, *Bnmfs*, was identified and mapped for its male phenotype by whole-genome resequencing and RNA-sequencing techniques (Teng et al., 2017). In another RGMS line, 117AB, inheritance analysis identified two genes involved in the genetic regulation of male sterility traits (Hou et al., 1990). In addition to these systems, several RGMS lines were controlled by three genes, such as: S45AB was regulated by *Bnms1-3* (He et al., 2008; Lei et al., 2007; Yi et al., 2006), 9012AB by *ms3-4* and its recessive epistatic suppressor gene (*esp*) (Ke et al., 2004; Xu et al., 2009) and RG206AB by two male sterility alleles (*BnRf* and *BnRf^b^*) and the restorer allele *BnRf*^a^ (Deng et al., 2016).

RGMS is characterized by the failure to produce functional anthers and pollen grains at various stages of anther microsporogenesis and dehiscence after cell division, differentiation, and subsequent degeneration (Goldberg et al., 1993; McCormick, 1993; Scott et al., 1991; Ying et al., 2003; Zhao et al., 2002b). The abnormal anthers have been programmed to form a four-lobed structure consisting of highly specialized reproductive tissues for pollen production and non-reproductive tissues (including epidermis, endothecium, middle layer and tapetum) for normal pollen development and the release of pollen grain release (Goldberg et al., 1993; Zhao et al., 2002a). The tapetum is the major tissue provides precursors for pollen development and pollen wall formation, and is highly regulated by several reported genes, including *DYSFUNCTIONAL TAPETUM1*(*DYT1*), *DEFECTIVE in TAPETAL DEVELOPMENT, FUNCTION1* (*TDF1*), *ABORTED MICROSPORES* (*AMS*), and *AtMYB103*/*MS188* (Chen et al., 2009). In addition, the whole process of pollen development and pollination plays a key role in the male fertility of flowering plants, and this process could be controlled by *CALLOSE SYNTHASE-LIKE* genes or *GLUCAN SYNTHASE-LIKE* genes, *ARABINOGALACTAN PROTEIN 11* (*AGP11*), *REVERSIBLY GLYCOSYLATED POLYPEPTIDE 1* and *2* (*RGP1* and *2*), and *CYSTEINE ENDOPEPTIDASE 1* (*CEP1*) (Chen et al., 2009; Dong et al., 2005; Ellinger, 2014; Enns et al., 2005; Huang et al., 2009; Nishikawa et al., 2005; Töller et al., 2008; Xie et al., 2012a; Xu et al., 2016; Záveská Drábková and Honys, 2017; Zhang et al., 2014) and so on. The mutation of these key genes might result in the appearance of a male sterility phenotype.

Apart from the traditional techniques for genetic research, such as QTL mapping, numerous novel techniques based on sequencing have emerged in recent years. The sequencing-based bulked segregant analysis (BSA-Seq) strategy was a more efficient QTL mapping method compared to the traditional QTL mapping method. By combining the traditional BSA method and the next-generation sequencing (NGS) technique, the BSA-Seq technique is widely used in QTL mapping, for example in rapeseed (Jia et al., 2023; Nan et al., 2024), rice (Gao et al., 2023), corn (Zhou et al., 2023) and so on. Sequence analysis of the QTL region by BSA-Seq could also identify many single nucleotide polymorphisms (SNPs) and small insertions and deletions (InDels). The specific markers could be designed and used for fine mapping of the candidate gene on the primary QTL regions and identified lots of genes (Liu et al., 2020). In the present study, the mechanism of a new RGMS line of 1205A was investigated by fine mapping and qRT-PCR analysis. Metabolic variation caused by the male sterility gene was also examined. The study renews the annotation of the gene and provides new resources for basic research on the genetic control of male sterility.

## Materials and Methods

### Plant materials and field experiment

The RGMS line of 1205AB was developed from a spontaneous mutant in 2010 by the Oil Crops Research Group of Guizhou University, Guiyang, Guizhou, China. The steps to establish the segregating population with sterile and fertile plants are shown in Figure 1. 1205AB was developed and is used in our research group as a two-line system for rapeseed hybrid production. Three populations from the cross between 1205A and 1205B were planted a in 2016 (730 plants), 2017 (830 plants) and 2018 (1,424 plants) and examined for plant sterility. The plant without pollen from all flowers was recorded as sterile, otherwise fertile. Subsequently, in 2020, 1205A was crossed with a new-type *B. napus* line of NT7G132, which was developed from *B. carinata* and *B. rapa* (Tian et al., 2010; Xiao et al., 2010) in 2020. Twenty-three hybrid combinations were performed between sterile plants and fertile plants in the F_2_ population with 162 plants (45 sterile plants and 117 fertile plants). In F_3_, 7 hybrid combinations showed an exact 1:1 segregation, and a total of 173 plants were used for plant sterility testing. One of the seven hybrid combinations with more progenies was renamed as NT7G132AB*^Bna1205ams1^* for future fine mapping. A total of 1,847 F_4_ plants from NT7G132A*^Bna1205ams^*×NT7G132B*^Bna1205ams1^*were planted and used for sterility investigation and fine mapping. All plants were planted in the Teaching and Practice Base of Guizhou University (106.683651°N, 26.410435°W) in Guiyang, China. Each row had 30 plants with a plot size of 4 m × 0.4 m.

**Figure 1.**
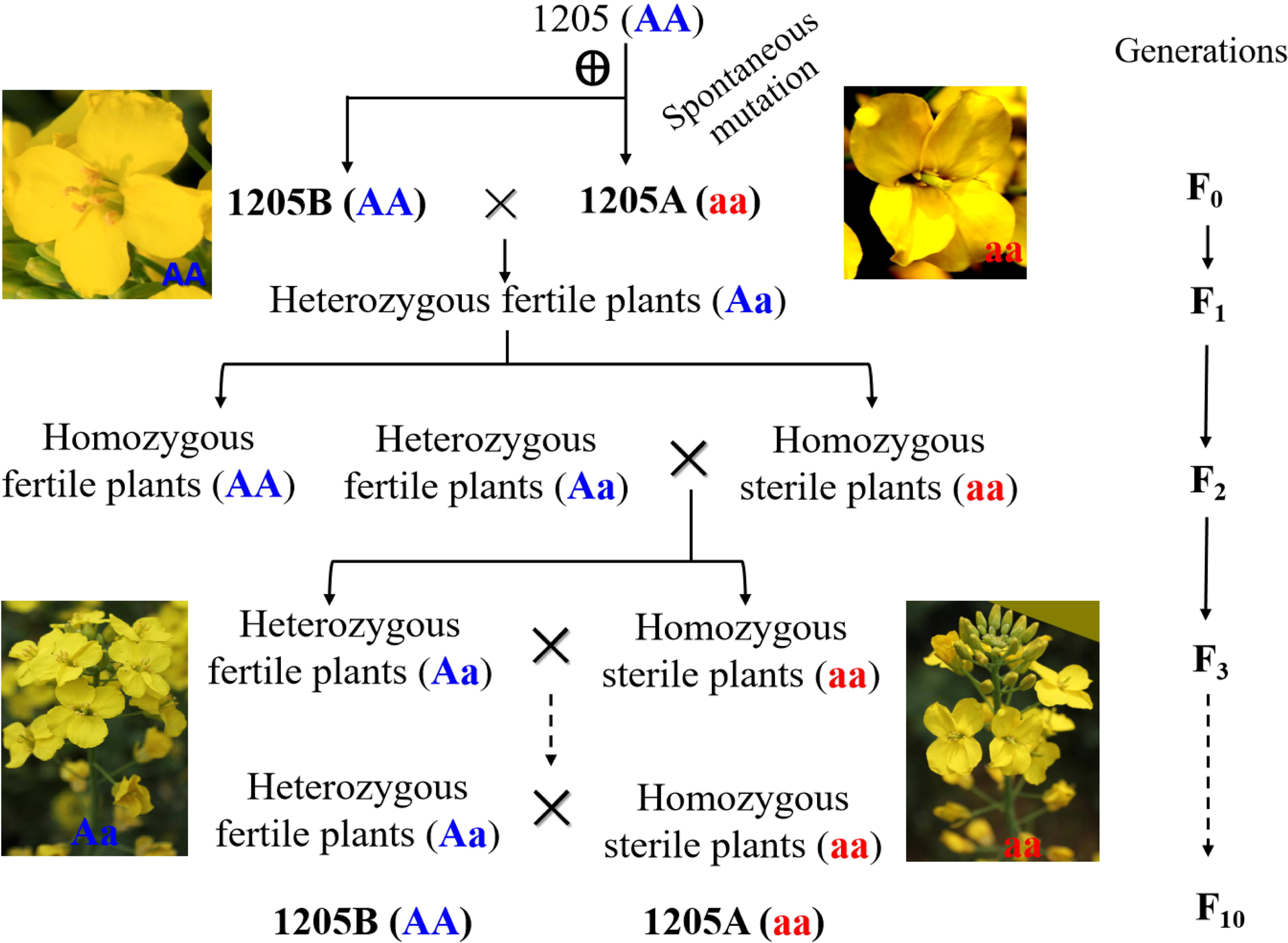
Schematic and workflow for the development of the 1205AB lines. The letter “A” in brackets indicates the fertile allele, compared to the recessive allele by a letter of “a” in parentheses. Genotypes “AA” and “Aa” were fertile and the genotype “aa” was sterile. 1205 indicates the line number.

### DNA library construction and BSA-Seq analysis

Total genomic DNA from the young leaves of 30 fertile plants and 30 sterile plants of 1205AB in 2018 was extracted using the cetyltrimethylammonium bromide (CTAB) method (Paterson et al., 1993) with minor modifications. Equal proportions of DNA from 30 fertile plants and 30 sterile plants were mixed and processed into a fertile pool (FP) and a sterile pool (SP), respectively. The sequencing library was constructed from 5 μg each of FP and DNA samples and sequenced in 76 cycles on an Illumina Novaseq 6000 platform using the PE150 sequencing model by LC-BIO TECHNOLOGIES CO., LTD. (www.lc-bio.com, Hangzhou, China). Adaptor reads, reads with >5% undetermined nucleotides, low quality (Q≤10) nucleotides occupying more than 20% were trimmed. The high-quality clean reads were then aligned to the *B. napus* reference genome of “ Darmor-bzh” (https://www.genoscope.cns.fr/brassicanapus/) (Chalhoub et al., 2014) using the Burrows–Wheeler Aligner software (BWA; version 0.7.13) (Li and Durbin, 2010). Single nucleotide polymorphisms (SNPs) and insertions and deletions (InDels) were detected across FP and SP by using GATK software version 3.8.1 (McKenna et al., 2010). Then, the ED (Euclidean distance) method was used to analyze the candidate region between FP and SP. The parameters of ED were calculated as follows:

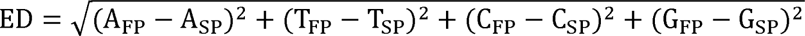

In the above equation, A, T, C, and G are the four base types. Sliding window analysis was used to display the distribution of ED values along the 19 *B. napus* chromosomes within 1 Mb width windows and 1 kb at each step using an in-house-developed Python script (available on request) based on the developed program (Takagi et al., 2013). The larger the ED value, the more likely it is that the SNPs and InDels contribute to the male sterility trait or are linked to a gene that controls the trait.

### Development and validation of KASP markers

BSA-Seq enabled the identification of numerous SNPs and InDels between FP and SP in the QTL region on chrC03 and chrC04. The sequence variation was used to develop Kompetitive Allele Specific PCR (KASP) markers based on the previously published high-quality *B. napus* reference genome “Darmor-bzh” (https://www.genoscope.cns.fr/brassicanapus/) (Chalhoub et al., 2014). Twenty-one markers for each of the QTL region of chrC03 and chrC04 were designed using the online software BatchPrimer3 v1.0 (https://probes.pw.usda.gov/batchprimer3/) with the parameters set to a minimum primer length of 15 bp and a maximum of 30 bp, and an annealing temperature of 59°C to 61°C, respectively, and synthesized Tsingke Biotech Co., Ltd (Beijing, China). Each KASP marker contains two allele-specific forward primers (F1 and F2) and a common reverse primer (R) and based on the evenly distributed flanking sequences around the variant position. The two allele-specific forward primers had the same primer sequence except for the targeting SNP at the 3’ end to distinguish the nucleotide differences of the SNPs. For F1 and F1, the blue fluorescent tag sequence of FAM (5’- GAAGGTGACCAAGTTCATGCT-3’) and the red fluorescent tag sequence of VIC (5’- GAAGGTCGGAGTCAACGGATT-3’) were added to their 5’ end, respectively. The established principles and procedures for genotyping identification of KASP assays were used (Ren et al., 2020). All 42 markers were used to amplify the 30 fertile plants and 30 sterile plants for BAS-Seq of 1205AB. The amplified sequences were also analyzed, and only five markers of ID04, ID07, ID10, ID21 and ID23 were identified on chrC03 and showed possible co-segregation with fertility in the 1205AB population. Then, the 5 markers were used to amplify the plants of NT7G132A*^Bna1205ams^*×NT7G132B*^Bna1205ams1^*in 2023 for fine mapping.

### RNA extraction and qRT-PCR

At flowering time, four sets of buds were sampled (X means bud size): X<1 mm (group S), 1≤X<2 mm (group SM), 2≤X< 3 mm (group M) and 3≤X<4 mm (group L). Total RNA (2 µg) was extracted from different stages of S, SM, M and L with three independent plants for each of the 1205A and 1205B lines using the TRIzol kit (Invitrogen, Carlsbad, CA) according to the manufacturer’s instructions. The RNA purity was checked using a Kaiao K5500® Spectrophotometer (Kaiao, Beijing, China), and the RNA integrity and concentration were assessed using the RNA Nano 6000 Assay Kit for the Bioanalyzer 2100 system (Agilent Technologies, CA, USA). Then, the cDNA was synthesized by reverse transcription using the PrimeScript RT Reagent Kit with gDNA Eraser (Takara, Dalian, China) according to the manufacturer’s instructions. Twenty-two candidate gene-specific primers for qRT-PCR were designed based on reference unigene sequences using Primer Premier 5.0. Real-time PCR was conducted using SsoAdvanced^TM^ Universal SYBR`R` Green Supermix (Hercules, CA) according to our previous research (Khattak et al. 2019).

The 2^−ΔΔCt^ algorithm was used to calculate the relative level of gene expression. The β*-actin* gene served as an internal control. All qRT-PCR was performed with three biological replicates and run on a Bio-Rad CFX96 Real Time System (Bio-Rad, Hercules, CA, USA).

### Untargeted metabolomic analysis

The flower buds (80 mg) of different sizes (S, SM, M and L) from 6 independent plants (replicates) for each set and line of 1205A and 1205B were immediately frozen in liquid nitrogen and ground into fine powder using a mortar and pestle. All 48 samples were then sent to APTBIO (https://www.aptbiotech.com/company) for analysis. The detailed analytical steps were as follows: 1000 μl of methanol/acetonitrile/H2O (2:2:1, v/v/v) was added to the homogenized solution for metabolite extraction. The mixture was centrifuged (14000g, 4 °C) for 20 min. The supernatant was dried in a vacuum centrifuge. For LC-MS analysis, samples were redissolved in 100 μl of acetonitrile/water (1:1, v/v) solvent and centrifuged at 14000 g for 15 min at 4 LJ, then the supernatant was injected. Analyzes were performed using a UHPLC (1290 Infinity LC, Agilent Technologies) connected to a quadrupole time-of-flight (AB Sciex TripleTOF 6600) from Applied Protein Technology Co., Ltd (Shanghai, China).

The ESI source conditions were set as follows: ion source gas1 (Gas1) as 60, ion source gas2 69(Gas2) as 60, curtain gas (CUR) as 30, source temperature: 600LJ, ionspray voltage floating (ISVF)±5500 V. In MS only acquisition, the instrument was set to acquire over the m/z range 60-1000 Da, and the accumulation time for TOF MS scan was set at 0.20 s/spectra. In auto MS/MS acquisition, the instrument was set to acquire over the m/z range 25-1000 Da, and the accumulation time for the product ion scan was set to 0.05 s/spectra. The product ion scan is acquired using information dependent acquisition (IDA) with high sensitivity mode selected. The parameters were set as follows: the collision energy (CE) was set at 35 V with ± 15 eV; declustering potential (DP), 60 V (+) and −60 V (−); exclude isotopes within 4 Da, candidate ions to be monitored per cycle: 10.

The raw MS data (wiff.scan files) were converted to MzXML files using ProteoWizard MSConvert before importing into the freely available XCMS software. The following parameters were used for peak picking: centWave m/z = 10 ppm, peakwidth = c (10, 60), prefilter = c (10, 100). For peak grouping, bw = 5, mzwid = 0.025, minfrac = 0.5 were used. CAMERA (Collection of Algorithms of MEtabolite pRofile Annotation) was sued for annotating isotopes and adducts. Only those variables that had more than 50% of the nonzero readings in at least one group were retained in the extracted ion features. Identification of metabolite compounds was performed by comparing the accuracy of m/z value (<10 ppm) and MS/MS spectra with an in-house database created with available authentic standards. After normalization to total peak intensity, the processed data were analyzed by using the R package (ropls) and was subjected to multivariate data analysis, including Pareto-scaled principal component analysis (PCA) and orthogonal partial least-squares discriminant analysis (OPLS-DA). 7-fold cross-validation and response permutation tests were used to assess the robustness of the model. The variable importance in projection value (VIP) of each variable in the OPLS-DA model was calculated to indicate its contribution to the classification.

### Differential metabolites (DMs) analysis

The identified metabolites were compared between SP and FP at each stage of S, SM, M and L. Metabolites with a VIP value >1 were further applied to Student’s t-test at the univariate level to measure the significance of each metabolite, the p values below 0.05 were considered as statistically significant. The Mantel test between gene expression level and metabolic abundance in the four stages of S, SM, M and L was calculated in R package ggcor, while the visualized correlation heatmap was performed by chiplot (Ji et al., 2022).

## Results

### Male sterility of 1205A is controlled by one gene locus

The recessive genic male-sterile (RGMS) line of 1205AB was developed by our laboratory and used for hybrid production, in which any *B. napus* breeding line could restore its fertility (Fig. 1) (Ye, 2020). Consistent with other reported RGMS lines, male sterility of 1205A was stable, complete, and unaffected by the environment (Fig. 1). The segregation ratio of fertile/sterile plants in three 1205A×1205B-derived populations were examined. The number of fertile/sterile plants in the three populations was 356/374 in 2016, 402/428 in 2017 and 705/719 in 2018, fitting a segregation ratio of 1:1 (χ2 = 0.444, p = 0.505; χ2 = 0.138, p = 0.711; χ2 = 0.814, p = 0.367, respectively), suggesting that male sterility of 1205A was controlled by a gene locus in these 1205A×1205B-derived populations. The locus identified through the inheritance analysis was named *Bna1205ams1*. In addition, the gene locus of *Bna1205ams1* was transferred through backcrossing to a new-type *B. napus* line of NT7G132 developed from *B. carinata* and *B. rapa* (Tian et al., 2010; Xiao et al., 2010), creating a new RGMS line of NT7G132AB*^Bna1205ams1^*.

The segregation ratio of fertile/sterile plants in two populations derived from NT7G132A*^Bna1205ams^*×1205B NT7G132B*^Bna1205ams1^*were also examined. The number of fertile/sterile plants in the two populations was 91/82 in F_3_ in 2022, 956/891 in F_4_ in 2023, fitting a segregation ratio of 1:1 (χ2 = 0.468, p = 0.494; χ2 = 2.287, p = 0.130), further confirming that male sterility of 1205A was controlled by a gene locus of *Bna1205ams1*.

### Male sterility gene locus were mapped by BAS-Seq

To quickly map the gene locus of *Bna1205ams1*, the BSA-Seq strategy method was used. Thirty randomly selected plants from each of 1205A and 1205B were used for DNA extraction, equal amounts of which were mixed to form a sterile pool (SP) and a fertile pool (FP) for whole-genome resequencing. After filtering the raw data, 32.45 Gb of clean data were obtained for FP with a Q30% of 94.36% and 36.28 Gb of clean data for SP with a Q30% of 94.10%. Among these clean reads, a total of 210,153,972 reads (97.13%) for FP and 235,697,816 (97.46%) for SP were mapped to the reference genome of “Darmor-bzhof” in *B. napus* (Chalhoub et al., 2014). The sequencing depth of coverage with >20× was 43.84% for FP and 62.43% for SP (Fig. S1). A total of 6,706,272 SNPs were detected between FP and SP, including 54.84% of “intergenic_region” type, 11.62% of “upstream_gene_variant” type, 11.45% of “downstream_gene_variant” type (Table S1). In addition, a total of 1,155,082 insertions and deletions (InDels), including 49.48% of the “intergenic_region” type, 15.65% of the “upstream_gene_variant” type, 13.87% of the “intron_variant” type, 13.90% of the “downstream_gene_variant” type were detected between FP and SP (Table S2).

The SNP/InDel-index between FP and SP was calculated using the Euclidean distance algorithm (ED) and used for QTL mapping (Fig. 2; Fig.3A). Finally, two candidate QTL regions on chrC03 and chrC04 were obtained for the male sterility gene locus of *Bna1205ams1*. The first QTL on chrC03 spanned 3.54 Mb (15.36–18.90 Mb) (Fig. 2; Fig.3A) and included 31,751 SNPs, 4,684 InDels, and 416 candidate genes compared to the *B. napus* reference genome of “Darmor-bzh”. The second QTL on chrC04 spanned 2.85 Mb (2.43–5.28 Mb) (Fig. 2) and contained 32,117 SNPs, 4,979 InDels, and 575 candidate genes compared to the *B. napus* reference genome of “Darmor-bzh”.

**Figure 2.**
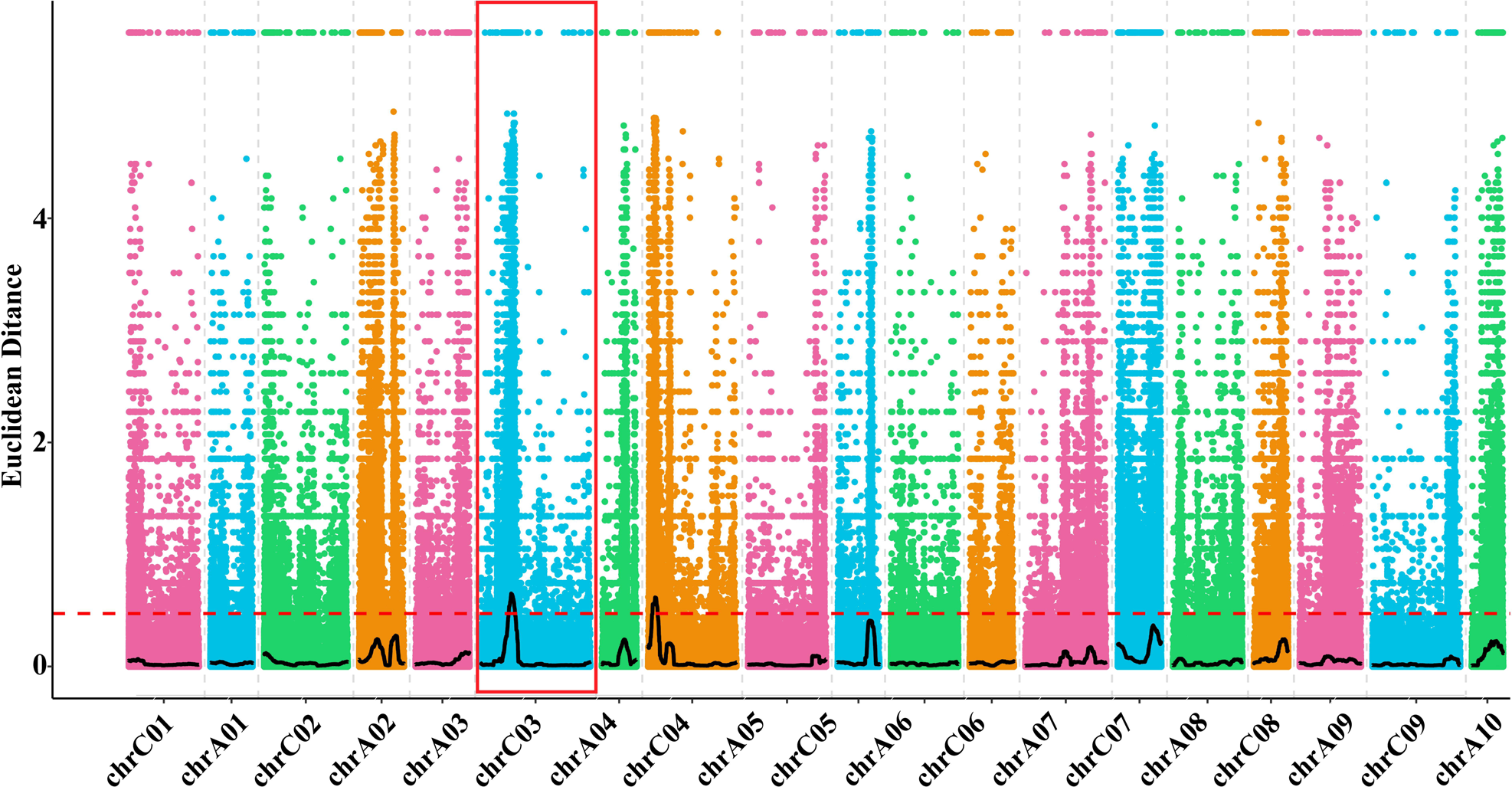
QTL mapping results by BSA-Seq method. The dashed red line indicates the threshold for QTL detection. chrC01 to chrC9 and chrA01-chrA10 on the x-axis indicate the 19 chromosomes of *B. napus* genome. The number on the y-axis indicates the values calculated by the Euclidean Distance algorithm.

**Figure 3.**
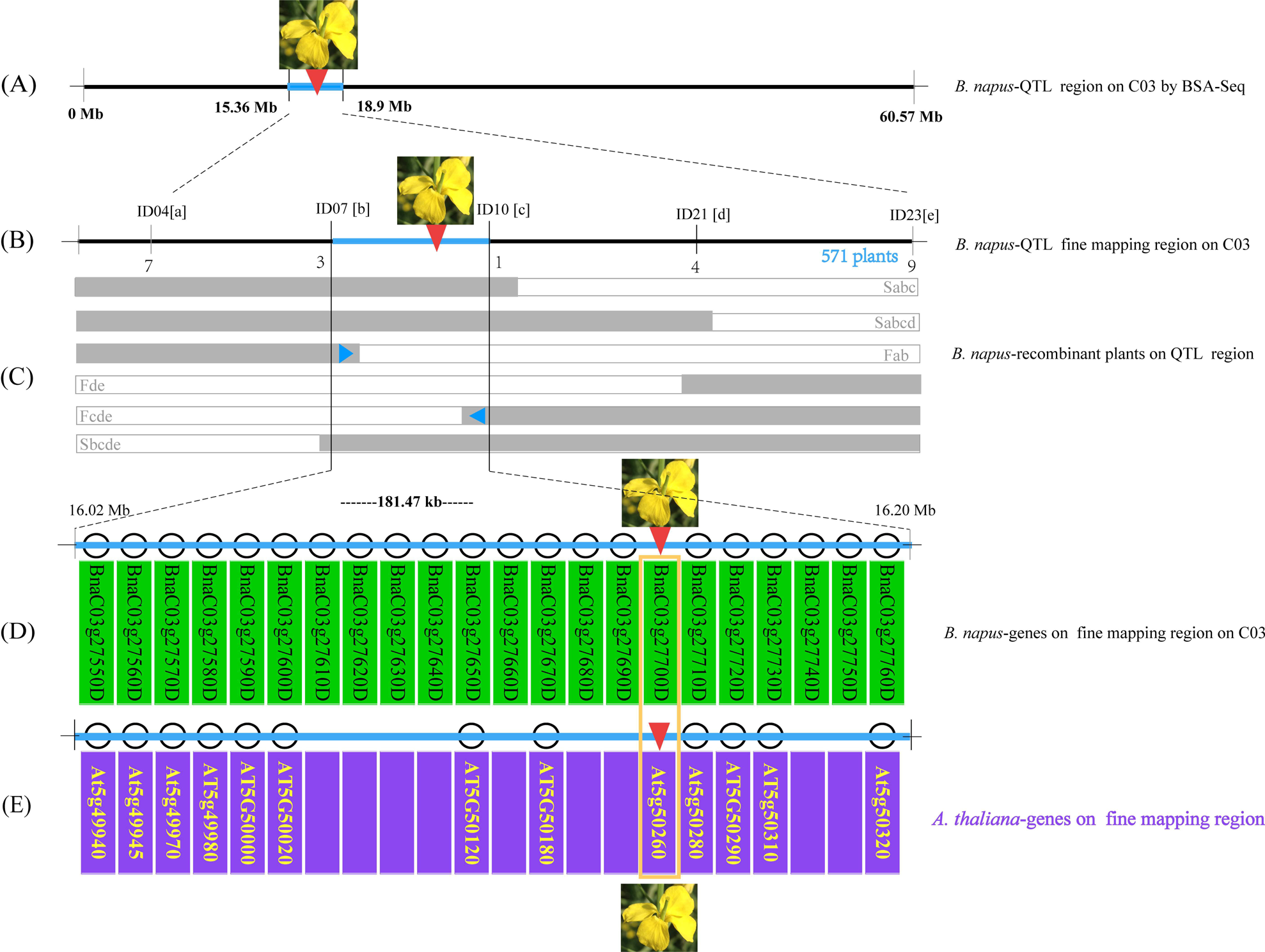
Fine mapping of the male sterility gene. (A) indicate the QTL mapping results by BSA-Seq. (B) and (C) indicate the results of the fine mapping, the numbers blow after C03 indicate the number of recombinants. ID04, ID07, ID10, ID21 and ID23 in the up of chrC03 indicate the five SNP markers developed in this study in (B). The letters “S” and “F” indicate the sterile plants and fertile pants, while the letters “a”, “b”, “c”, “d”, “e” indicate the recombinant type by the marker was recognizable detectable, for example, “Sabc” indicates that the recombinant plant was sterile and cloud be recognized by the makers of ID04 by “a”, ID07 by “b”, ID10 by “c”. “571” indicates the number of plants used for traditional fine mapping in (C). The twenty-two candidate genes based on the *B. napus* reference genome of “Darmor-bzh” in the fine-mapped QTL region were displayed in the 181.47 kb region under the chromosome (D). The sequence fine-mapped QTL region were compared with the Arabidopsis genome, and the corresponding allele in Arabidopsis were marked under chromosome region (E).

### Male sterility gene locus was validated and fine mapped

For fine mapping of *Bna1205ams1*, 21 kompetitive allele specific PCR (KASP) markers were developed based on the SNP and InDel variants, which were evenly distributed on each of the QTL regions of chrC03 and chrC04. The 42 SNP markers were further validated by amplification of genomic DNA of 30 fertile plants and 30 sterile plants once used in BSA-Seq analysis of 1205AB. Only five markers of ID04, ID07, ID10, ID21 and ID23 on chrC03 (Table S3) showed better amplification results and possible co-segregation with the male sterility trait of the 1205AB population. For fine mapping of *Bna1205ams1*, 571 F_4_ plants (285 fertile plants and 286 sterile plants) from the newly developed NT7G132AB*^Bna1205ams1^* population were used. After detecting the genotype of all these 571 F_4_ plants, 16 recombinant plants including all these five markers were detected (Fig. 3B). All 16 recombinants could be divided into the 6 genotypes of Sabc, Sabcd, Fab, Fde, Fcde and Sbcde (Fig. 3B). The fertility of the 16 recombinants was consistent with the genotype results. Finally, the QTL region including *Bna1205ams1* on chrC03 was narrowed down to 181.47 kb between the genotype of Fab and Fcde (Fig. 3C). All genes within the fine mapping region were analyzed, with 22 genes found compared to the *B. napus* reference genome of “Darmor-bzh” (Fig. 3D).

The sequence of the QTL region containing *Bna1205ams1* was compared to the Arabidopsis genome, which showed high sequence similarity to a fragment on chr05 between *At5g4940* and *At5g50320* of *A. thaliana* (Fig. 3D and 3E). The sequence of the 22 genes in the fine mapping region of chrC03 was blasted against the Arabidopsis genome in the TAIR database (https://www.arabidopsis.org/index.jsp), and 7 genes found their Arabidopsis alleles (Fig. 3D and 3E).

### The candidate genes for male sterility were identified

To further investigate the candidate genes of *Bna1205ams1*, the expression levels of the 22 genes in the fine mapping region of chrC03 at four different developmental stages of S, SM, M and L of flower buds were analyzed by qRT-PCR. Finally, only three genes of *BnaC03g27600D*, *BnaC03g27670D* and *BnaC03g27700D* showed significant differences on at least one stage (Table S3, Fig. 4). The expression of *BnaC03g27600D* showed a significant difference in L stage (p = 0.01) (Fig. 4A). *BnaC03g27670D* showed significant expression differences in SM (p = 0.05) and L stages (p = 0.001) (Fig. 4B). Furthermore, the expression level of *BnaC03g27700D* in the three stages of SM, M and L was significantly higher in 1205A than in 1205B (p = 0.001) (Fig. 4C). The large expression difference of *BnaC03g27700D* at different stages was also shown with a heatmap (Fig. 4D).

**Figure 4.**
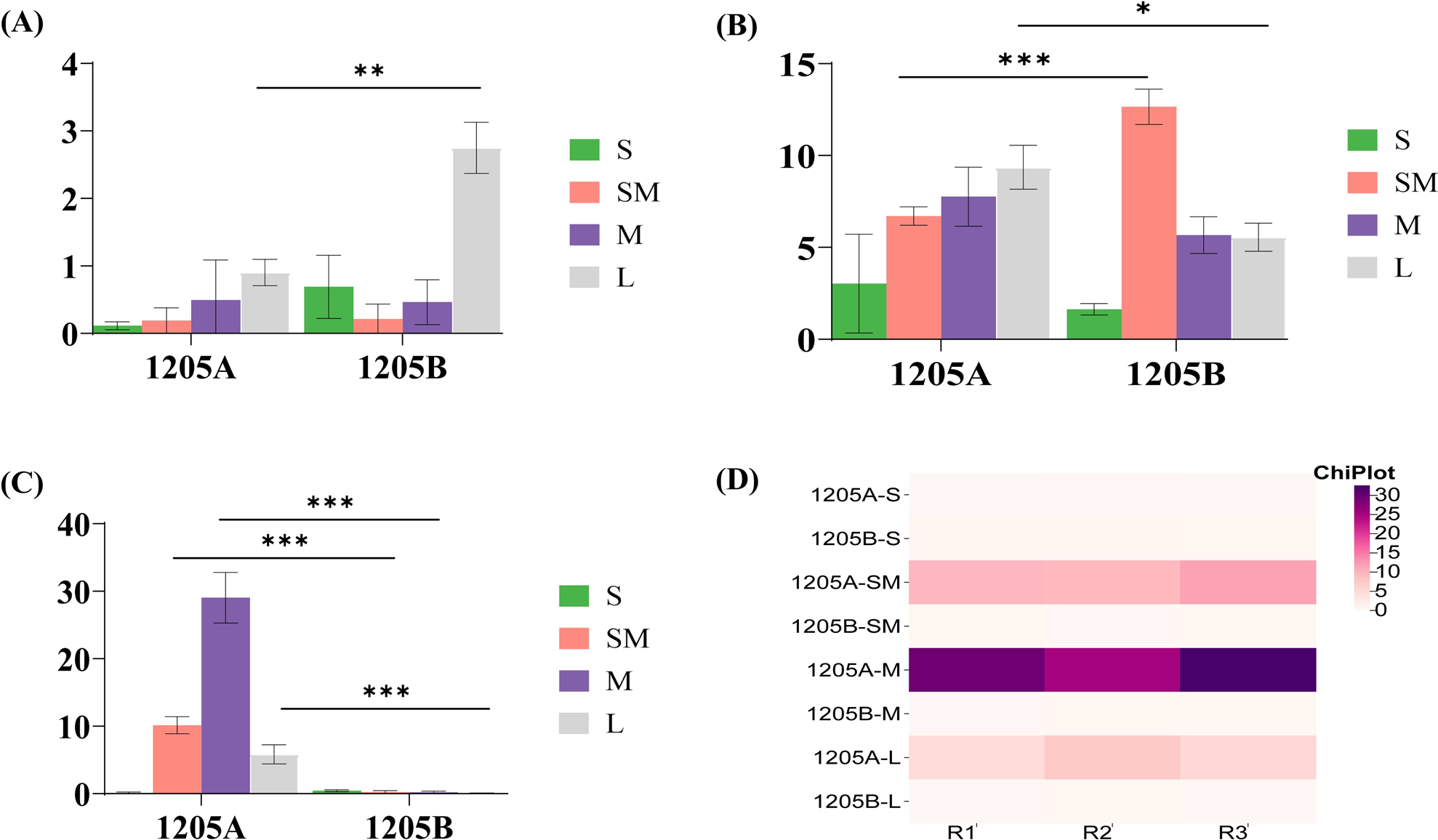
qRT-PCR analysis of the candidate gene in the fine-mapped QTL region. The genes with no difference in any of the four stages of S, SM, M and L between 1205A and 1205B were not shown. (A), (B), and (C) indicate the expression level of *BnaC03g27600D*, *BnaC03g27670D*, and *BnaC03g27700D*, respectively. The expression level of *BnaC03g27700D* was specifically analyzed using a heatmap, R_1_, R_2_ and R_3_ indicate three replicates. *, ** and *** indicate the significant difference at levels of p=0.05, p=0.01 and p=0.001.

The functions were analyzed for these three candidate genes. It has been reported that *At5g50020*, which corresponds to *Protein S-acyltransferase 9* (*PAT9*), is involved in plant immune response as an Arabidopsis allele of *BnaC03g27600D*, and its mutation could lead to increased plant immune response (Chen et al., 2019). *At5g50180* corresponds to *BnaC03g27670D*. It was predicted to be a member of mitogen-activated protein kinase (MAPK) and clustered with *ATN1*, which was predicted to be a signal transduction module (Group, 2002). While the exact function of *At5g50180* was not yet known from the published data. In addition, the Arabidopsis allele of *BnaC03g27700D*, *At5g50260* (*CEP1*, *CYSTEINE ENDOPEPTIDASE 1*) has been reported to be an important executor involved in tapetal programmed cell death and regulation of pollen development in Arabidopsis (Zhang et al., 2014). Thus, *BnaC03g27700D* showed a high probability of genetic regulation of male sterility of 1205A (Fig. 3F).

### Large metabolic fluctuations were caused by the male sterility gene mutation

To better explore and annotate the functions and consequences resulting from the *Bna1205ams1* mutation, metabolic analysis was performed in the four stages of S, SM, M and L in 1205A compared to 1205B. The metabolites of 48 samples, from four sets each of 1205A and 1205B with six replicates were analyzed. A total of 784 and 569 metabolites (Table S4) were identified using the positive ion mode (POS) and negative ion mode (NEG) methods, respectively. The metabolic level between 1205A and 1205B at the same stage was compared and a total of 168 differential metabolites (DMs) were identified. These DMs include 18 in S (15 down, 3 up), 39 in SM (19 down, 20 up), 67 in M (31 down, 36 up), 44 in L (26 down, 18 up) (Fig. 5A). The function of DMs was analyzed by KEGG (Kyoto Encyclopedia of Genes and Genomes). In the S stage, the first three functional classes of DMs, including 7 DMs for lipids and lipid-like molecules, 5 DMs for phenylpropanoids and polyketides, 3 DMs for each of organoheterocyclic compounds and nucleosides, nucleotides, and analogues (Fig. 5B). In the SM stage, the first three functional classes of DMs, including 18 DMs for lipids and lipid-like molecules, 8 DMs for phenylpropanoids and polyketides, and 8 DMs for organoheterocyclic compounds (Fig. 5B). In the M stage, the first three functional classes of DMs, including 18 DMs for lipids and lipid-like molecules, 14 DMs for organic acids and derivatives, and 13 DMs for phenylpropanoids and polyketides (Fig. 5B). In the L stage, the first three functional classes of DMs, including 12 DMs for lipids and lipid-like molecules, 12 DMs for phenylpropanoids and polyketides, and 4 DMs for each of organic acids and derivatives and organoheterocyclic compounds (Fig. 5B).

**Figure 5.**
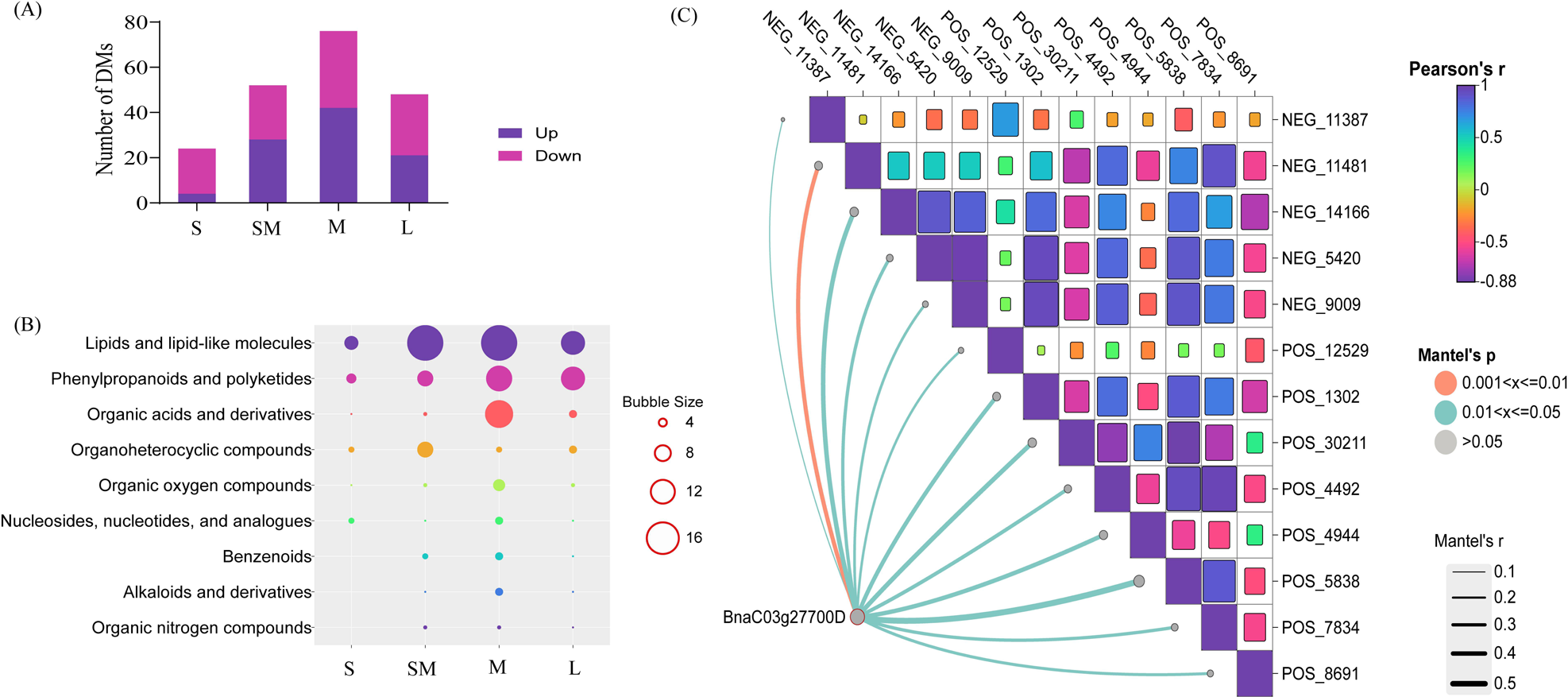
Analysis of the metabolic fluctuations resulted from the mutation of male sterility gene. The number of differential metabolites (DMs) between 1205A and 1205B at four stages of S, SM, M and L were analyzed in (A). The functional classes were analyzed for the DMs in the four stages of S, SM, M and L, the larger bubble size indicates more DMs in (B). The correlation heatmap between the expression level of *BnaC03g27700D* and the abundance of metabolites at four stages of S, SM, M and L was analyzed and shown using the Mantel test, while the correlation heatmap between different identified metabolites at four stages of S, SM, M and L was analyzed using the Pearson method in (C).

The correlation analysis between the expression level for the candidate gene of *BnaC03g27700D* in each of the four stages of S, SM, M and L and the corresponding abundance of each DM was performed. The expression level of *BnaC03g27700D* was significantly correlated with 13 metabolites, including four lipids and lipid-like molecules, three phenylpropanoids and polyketides, two organoheterocyclic compounds, one each for organic nitrogen compounds, organic oxygen compounds and nucleosides, nucleotides (Fig. 5C; Table S5).

## Discussion

In China, RGMS has become an important two-line hybrid system, and several RGMS-based lines have already been registered and used for hybrid production (FX et al., 1993; Li et al., 2012a; Yi et al., 2006). Compared with the three-line hybrid system, the two-line hybrid system has the following characteristics: complete and stable male sterility, easy transfer of the sterility trait to different genetic backgrounds, abundant recovery resources (Lei et al., 2007), and simplified hybrid production system. These features of the two-line system are extremely attractive for regions with unstable temperatures at the flowering time of *B. napus*, such as some parts of Guizhou Province in China, where the temperature often fluctuates between 0°C and 10°C. This may allow the emergence of some fertile pollen in the CMS lines.

In the present study, the male sterility regulation mechanism of 1205A was investigated. As an RGMS line for two-line hybrid production, 1205A was demonstrated to be controlled by one gene locus in this study and mapped on chrC03 by BSA-Seq and linkage mapping. Furthermore, the QTL region was narrowed down to a 181.47 kb region by fine mapping. Through expression and functional analysis of 22 genes in the fine-mapping QTL region, *BnaC03g27700D* showed a higher probability as a candidate gene for the genetic regulation of male sterility in 1205A. *At5g50260* (*CEP1*, *CYSTEINE ENDOPEPTIDASE 1*) was the corresponding allele of *BnaC03g27700D* in *A. thaliana*, which had been reported to be involved in tapetal programmed cell death and regulation of pollen development in Arabidopsis (Zhang et al., 2014).

The functions of *BnaC03g27700D* were further validated by metabolic analysis between 1205A and 1205B. The expression level of *BnaC03g27700D* was consistent with the number of DMs in the present study. In detail, the metabolic pathways identified many DMs, most of which were involved in aborted tapetal PCD, which could lead to reduced pollen fertility with abnormal pollen exine (Xu et al., 2022; Zhang et al., 2014). Through correlation analysis, 13 metabolites were significantly correlated with the expression level at each stage of S, SM, M and L, which provided further potential functions for *BnaC03g27700D* in our future study.

Different from the regulatory model in Arabidopsis, the candidate gene of *BnaC03g27700D* was almost not expressed in the fertile plants of 1205B and was extremely highly expressed in 1205A from the SM stage to the L stage. From this, we infer that there may be an epistatic suppressor gene in 1205B that restricts the expression of *BnaC03g27700D*. Upon loss of the epistatic suppressor gene in 1205A, *BnaC03g27700D* expressed highly and resulted in abnormal PCD and pollen development. The epistatic suppressor gene was widespread in Brassiceae. A tri-genetic inheritance model, later proposed to interpret sterility inheritance in 9012AB, suggests that male sterility is controlled by the interaction of two recessive male-sterile genes (*Bnms3* and *Bnms4*) with an epistatic suppressor gene (*BnRf* or *BnEsp*) (Chen et al., 1998). In other reports on *B. napus* (Song et al., 2006) and *B. rapa* (Feng et al., 2009), the epistatic suppressor gene was also found to be the same genetic pattern. Compared to other reported two-line hybrid breeding system, to our knowledge, line 1205AB should be a new one because the male sterility genes in *BnaC03g27700D* were completely different, e.g., *BnMs3* on C09 (Li et al., 2012a), *BnRf* on A07 (Xie et al., 2012b) in 9012AB, *Bnms1* on A07 (Yi et al., 2006) and *BnMs2* on C06 in S45AB (Yi et al., 2010).

In summary, the study reported a new two-line hybrid production system for 1205AB and a newly constructed two-line hybrid production system for NT7G132A*^Bna1205ams^*. The new RGMS line 1205A provided us with the opportunity to identify a new male sterility gene of *BnaC03g27700D* in *B. napus*. The study of *BnaC03g27700D* aims to renew the annotation of the gene and provide new resources for basic research on the genetic control of male sterility.

## Author Contributions

ET conceived and designed the experiments. LX and JZ performed the experiments. LX, JZ and ET analyzed the data and wrote the paper. LX, JZ, KY, XY, QW, HJ and QO revised the paper.

## Compliance with ethical standards

### Ethical standards

The experiments were performed in compliance with the current laws of China.

## Supporting information

**Figure S1 Sequencing depth of coverage in BSA-Seq analysis.**

**Table S1 Information on SNP variation detected in BSA-Seq analysis.**

**Table S2 Information on InDel variation detected in BSA-Seq analysis.**

**Table S3 Primers used in this study.**

**Table S4 The differential metabolites detected in this study.**

**Table S5 The detailed information of the 13 metabolites correlates closely with the candidate gene.**

## References

Chalhoub, B., Denoeud, F., Liu, S., Parkin, I. A. P., Tang, H., Wang, X., Chiquet, J., Belcram, H., Tong, C., Samans, B. et al. (2014). Early allopolyploid evolution in the post-Neolithic Brassica napus oilseed genome. Science 345, 950–953.

Chen, D., Ahsan, N., Thelen, J. J. and Stacey, G. (2019). S-Acylation of plant immune receptors mediates immune signaling in plasma membrane nanodomains. BioRxiv, 720482.

Chen, F., Hu, B., Li, C., Li, Q., Chen, W. and Zhang, M. (1998). Genetic studies on GMS in Brassica napus LI Inheritance of recessive GMS line 9012A.

Chen, Y., Lei, S., Zhou, Z., Zeng, F., Yi, B., Wen, J., Shen, J., Ma, C., Tu, J. and Fu, T. (2009). Analysis of gene expression profile in pollen development of recessive genic male sterile Brassica napus L. line S45A. Plant Cell Reports 28, 1363–1372.

Cheng, Q., Wang, P., Liu, J., Wu, L., Zhang, Z., Li, T., Gao, W., Yang, W., Sun, L. and Shen, H. (2018). Identification of candidate genes underlying genic male-sterile msc-1 locus via genome resequencing in Capsicum annuum L. Theoretical and Applied Genetics 131, 1861–1872.

Deng, Z., Li, X., Wang, Z., Jiang, Y., Wan, L., Dong, F., Chen, F., Hong, D. and Yang, G. (2016). Map-based cloning reveals the complex organization of the BnRf locus and leads to the identification of BnRf b, a male sterility gene, in Brassica napus. Theoretical and Applied Genetics 129, 53–64.

Dong, X., Hong, Z., Sivaramakrishnan, M., Mahfouz, M. and Verma, D. P. S. (2005). Callose synthase (CalS5) is required for exine formation during microgametogenesis and for pollen viability in Arabidopsis. The Plant Journal 42, 315–328.

Ellinger, D. (2014). Callose biosynthesis in arabidopsis with a focus on pathogen response: what we have learned within the last decade. Annals of botany v. 114, pp. 1349–1358-2014 v.114 no.6.

Enns, L. C., Kanaoka, M. M., Torii, K. U., Comai, L., Okada, K. and Cleland, R. E. (2005). Two callose synthases, GSL1 and GSL5, play an essential and redundant role in plant and pollen development and in fertility. Plant Molecular Biology 58, 333–349.

Feng, H., Wei, P., Piao, Z.-Y., Liu, Z.-Y., Li, C.-Y., Wang, Y.-G., Ji, R.-Q., Ji, S.-J., Zou, T. and Choi, S.-R. (2009). SSR and SCAR mapping of a multiple-allele male-sterile gene in Chinese cabbage (Brassica rapa L.). Theoretical and Applied Genetics 119, 333–339.

Fu, T. (2009). Considerations on heterosis utilization in rapeseed (Brassica napus L.). 16th Australian research assembly on Brassicas, Ballarat.

FX, C., BC, H. and QS, L. (1993). Discovery and study of genic male sterility (GMS) material 9012A in Brassica napus L. Acta Agric Univ Pekinensis 19(Suppl).

Gao, Q., Wang, H., Yin, X., Wang, F., Hu, S., Liu, W., Chen, L., Dai, X. and Liang, M. (2023). Identification of Salt Tolerance Related Candidate Genes in ‘Sea Rice 86’at the Seedling and Reproductive Stages Using QTL-Seq and BSA-Seq. Genes 14, 458.

Goldberg, R. B., Beals, T. P. and Sanders, P. M. (1993). Anther development: basic principles and practical applications. The Plant Cell 5, 1217–1229.

Group, M. (2002). Mitogen-activated protein kinase cascades in plants: a new nomenclature. Trends in Plant Science 7, 301–308.

Guo, J., Zhang, Y., Hui, M., Cheng, Y., Zhang, E. and Xu, Z. (2016). Transcriptome sequencing and de novo analysis of a recessive genic male sterile line in cabbage (Brassica oleracea L. var. capitata). Molecular Breeding 36, 117.

Gupta, S. (2007). Advances in Botanical Research: Rapeseed Breeding. Elsevier, 45.

Han, Y., Zhao, F., Gao, S., Wang, X., Wei, A., Chen, Z., Liu, N., Tong, X., Fu, X., Wen, C. et al. (2018). Fine mapping of a male sterility gene ms-3 in a novel cucumber (Cucumis sativus L.) mutant. Theoretical and Applied Genetics 131, 449–460.

He, J., Ke, L., Hong, D., Xie, Y., Wang, G., Liu, P. and Yang, G. (2008). Fine mapping of a recessive genic male sterility gene (Bnms3) in rapeseed (Brassica napus) with AFLP- and Arabidopsis-derived PCR markers. Theoretical and Applied Genetics 117, 11–18.

Hou, G. Z., Wang, H. and Zhang, R. M. (1990). Genetic study on genic male sterility (GMS) material No. 117A in Brassica napus L. Oil Crop China 2.

Hu, J. Q., Wang, L. L., Xiang, T. H. and Pang, J. L. (2008). [Studies on the megasporogenesis and microsporegenesis and the development of the female gametophyte and male gametophyte in Camptotheca acuminate]. Fen Zi Xi Bao Sheng Wu Xue Bao 41, 367–75.

Huang, L., Chen, X.-Y., Rim, Y., Han, X., Cho, W. K., Kim, S.-W. and Kim, J.-Y. (2009). Arabidopsis glucan synthase-like 10 functions in male gametogenesis. Journal of Plant Physiology 166, 344–352.

Ji, J.-l., Yang, L.-m., Fang, Z.-y., Zhuang, M., Zhang, Y.-y., Lv, H.-h., Liu, Y.-m. and Li, Z.-s. (2017). Recessive male sterility in cabbage (Brassica oleracea var. capitata) caused by loss of function of BoCYP704B1 due to the insertion of a LTR-retrotransposon. Theoretical and Applied Genetics 130, 1441–1451.

Ji, X., Tang, J. and Zhang, J. (2022). Effects of Salt Stress on the Morphology, Growth and Physiological Parameters of Juglansmicrocarpa L. Seedlings. Plants 11, 2381.

Jia, X., Wang, S., Zhao, H., Zhu, J., Li, M. and Wang, G. (2023). QTL mapping and BSA-seq map a major QTL for the node of the first fruiting branch in cotton. Frontiers in Plant Science 14, 1113059.

Ke, L., Sun, Y., Liu, P. and Yang, G. (2004). Identification of AFLP fragments linked to one recessive genic male sterility (RGMS) in rapeseed (Brassica napus L.) and conversion to SCAR markers for marker-aided selection. Euphytica 138, 163–168.

Lei, S., Yao, X., Yi, B., Chen, W., Ma, C., Tu, J. and Fu, T. (2007). Towards map-based cloning: fine mapping of a recessive genic male-sterile gene (BnMs2) in Brassica napus L. and syntenic region identification based on the Arabidopsis thaliana genome sequences. Theoretical and Applied Genetics 115, 643–651.

Li, H. and Durbin, R. (2010). Fast and accurate long-read alignment with Burrows–Wheeler transform. Bioinformatics 26, 589–595.

Li, J., Hong, D., He, J., Ma, L., Wan, L., Liu, P. and Yang, G. (2012a). Map-based cloning of a recessive genic male sterility locus in Brassicanapus L. and development of its functional marker. Theoretical and Applied Genetics 125, 223–234.

Li, Z. Y., Wang, Y., Shen, W. T. and Zhou, P. (2012b). Content determination of benzyl glucosinolate and anti-cancer activity of its hydrolysis product in Carica papaya L. Asian Pac J Trop Med 5, 231–3.

Liu, Y., Zhou, X., Yan, M., Wang, P., Wang, H., Xin, Q., Yang, L., Hong, D. and Yang, G. (2020). Fine mapping and candidate gene analysis of a seed glucosinolate content QTL, qGSL-C2, in rapeseed (Brassica napus L.). Theoretical and Applied Genetics 133, 479–490.

McCormick, S. (1993). Male Gametophyte Development. Plant Cell 5, 1265–1275.

McKenna, A., Hanna, M., Banks, E., Sivachenko, A., Cibulskis, K., Kernytsky, A., Garimella, K., Altshuler, D., Gabriel, S. and Daly, M. (2010). The Genome Analysis Toolkit: a MapReduce framework for analyzing next-generation DNA sequencing data. Genome Research 20, 1297–1303.

Mishima, K., Hirao, T., Tsubomura, M., Tamura, M., Kurita, M., Nose, M., Hanaoka, S., Takahashi, M. and Watanabe, A. (2018). Identification of novel putative causative genes and genetic marker for male sterility in Japanese cedar (Cryptomeria japonica D.Don). BMC Genomics 19, 277.

Nan, Y., Xie, Y., He, H., Wu, H., Gao, L., Atif, A., Zhang, Y., Tian, H., Hui, J. and Gao, Y. (2024). Integrated BSA-seq and RNA-seq analysis to identify candidate genes associated with nitrogen utilization efficiency (NUtE) in rapeseed (Brassica napus L.). International Journal of Biological Macromolecules 254, 127771.

Nishikawa, S.-i., Zinkl, G. M., Swanson, R. J., Maruyama, D. and Preuss, D. (2005). Callose (β-1,3 glucan) is essential for Arabidopsispollen wall patterning, but not tube growth. BMC Plant Biology 5, 22.

Paterson, A. H., Brubaker, C. L. and Wendel, J. F. (1993). A rapid method for extraction of cotton (Gossypium spp.) genomic DNA suitable for RFLP or PCR analysis. Plant Molecular Biology Reporter 11, 122–127.

Prakash, S., Kirti, P. B., Bhat, S. R., Gaikwad, K., Kumar, V. D. and Chopra, V. L. (1998). A Moricandia arvensis– based cytoplasmic male sterility and fertility restoration system in Brassica juncea. Theoretical and Applied Genetics 97, 488–492.

Raja, D., Kumar, M. S., Devi, P. R., SLoganathan, a., Ramya, K., Kannan, N. and Subramanian, V. (2018). Identification of molecular markers associated with genic male sterility in tetraploid cotton (Gossypium hirsutum L.) through bulk segregant analysis using a cotton SNP 63K array. Czech Journal of Genetics and Plant Breeding 197, 161–173.

Ren, R., Xu, J., Zhang, M., Liu, G., Yao, X., Zhu, L. and Hou, Q. (2020). Identification and molecular mapping of a gummy stem blight resistance gene in wild watermelon (Citrullus amarus) germplasm PI 189225. Plant Disease 104, 16–24.

Scott, R., Hodge, R., Paul, W. and Draper, J. (1991). The molecular biology of anther differentiation. Plant Science 80, 167–191.

Song, L.-Q., Fu, T.-D., Tu, J.-X., Ma, C.-Z. and Yang, G.-S. (2006). Molecular validation of multiple allele inheritance for dominant genic male sterility gene in Brassica napus L. Theoretical and Applied Genetics 113, 55–62.

Töller, A., Brownfield, L., Neu, C., Twell, D. and Schulze-Lefert, P. (2008). Dual function of Arabidopsis glucan synthase-like genes GSL8 and GSL10 in male gametophyte development and plant growth. The Plant Journal 54, 911–923.

Takagi, H., Abe, A., Yoshida, K., Kosugi, S., Natsume, S., Mitsuoka, C., Uemura, A., Utsushi, H., Tamiru, M. and Takuno, S. (2013). QTL-seq: rapid mapping of quantitative trait loci in rice by whole genome resequencing of DNA from two bulked populations. The Plant Journal 74, 174–183.

Teng, C., Du, D., Xiao, L., Yu, Q., Shang, G. and Zhao, Z. (2017). Mapping and Identifying a Candidate Gene (Bnmfs) for Female-Male Sterility through Whole-Genome Resequencing and RNA-Seq in Rapeseed (Brassica napus L.). Frontiers in Plant Science 8.

Tian, E., Jiang, Y., Chen, L., Zou, J., Liu, F. and Meng, J. (2010). Synthesis of a *Brassica* trigenomic allohexaploid (*B. carinata* × *B. rapa*) de novo and its stability in subsequent generations. Theoretical and Applied Genetics 121, 1431–1440.

Wang, P., Bian, L. and Cao, C. (2016). Mapping of Nuclear Male-Sterile Gene ms14 Using SSR Markers in Cotton. Molecular Plant Breeding 7.

Xiao, Y., Chen, L., Zou, J., Tian, E., Xia, W. and Meng, J. (2010). Development of a population for substantial new type Brassica napus diversified at both A/C genomes. Theoretical and Applied Genetics 121, 1141–1150.

Xie, B., Deng, Y., Kanaoka, M. M., Okada, K. and Hong, Z. (2012a). Expression of Arabidopsis callose synthase 5 results in callose accumulation and cell wall permeability alteration. Plant Science 183, 1–8.

Xie, Y., Dong, F., Hong, D., Wan, L., Liu, P. and Yang, G. (2012b). Exploiting comparative mapping among Brassica species to accelerate the physical delimitation of a genic male-sterile locus (BnRf) in Brassica napus. Theoretical and Applied Genetics 125, 211–222.

Xu, B., Wu, R., Shi, F., Gao, C. and Wang, J. (2022). Transcriptome profiling of flower buds of male-sterile lines provides new insights into male sterility mechanism in alfalfa. BMC Plant Biology 22, 199.

Xu, T., Zhang, C., Zhou, Q. and Yang, Z.-N. (2016). Pollen wall pattern in Arabidopsis. Science Bulletin 61, 832–837.

Xu, Z., Xie, Y., Hong, D., Liu, P. and Yang, G. (2009). Fine mapping of the epistatic suppressor gene (esp) of a recessive genic male sterility in rapeseed (Brassica napus L.). Genome 52, 755–760.

Ye, B. (2020). Mapping and screening of candidate genes for regulation of sterility of 1205AB in Brassica napus, vol. Master: Guizhou University.

Yi, B., Chen, Y., Lei, S., Tu, J. and Fu, T. (2006). Fine mapping of the recessive genic male-sterile gene (Bnms1) in Brassica napus L. Theoretical and Applied Genetics 113, 643–650.

Yi, B., Zeng, F., Lei, S., Chen, Y., Yao, X., Zhu, Y., Wen, J., Shen, J., Ma, C., Tu, J. et al. (2010). Two duplicate CYP704B1-homologous genes BnMs1 and BnMs2 are required for pollen exine formation and tapetal development in Brassica napus. The Plant Journal 63, 925–938.

Ying, M., Dreyer, F., Cai, D. and Jung, C. (2003). Molecular markers for genic male sterility in Chinese cabbage. Euphytica 132, 227–234.

Záveská Drábková, L. and Honys, D. (2017). Evolutionary history of callose synthases in terrestrial plants with emphasis on proteins involved in male gametophyte development. PLOS ONE 12, e0187331.

Zhang, D., Liu, D., Lv, X., Wang, Y., Xun, Z., Liu, Z., Li, F. and Lu, H. (2014). The cysteine protease CEP1, a key executor involved in tapetal programmed cell death, regulates pollen development in Arabidopsis. The Plant Cell 26, 2939–2961.

Zhao, D.-Z., Wang, G.-F., Speal, B. and Ma, H. (2002a). The EXCESS MICROSPOROCYTES1 gene encodes a putative leucine-rich repeat receptor protein kinase that controls somatic and reproductive cell fates in the Arabidopsis anther. Genes & Development 16, 2021–2031.

Zhao, Y., Hull, A. K., Gupta, N. R., Goss, K. A., Alonso, J., Ecker, J. R., Normanly, J., Chory, J. and Celenza, J. L. (2002b). Trp-dependent auxin biosynthesis in Arabidopsis: involvement of cytochrome P450s CYP79B2 and CYP79B3. Genes & development 16, 3100–3112.

Zhou, W.-q., Zhang, H.-t., He, H.-j., Gong, D.-m., Yang, Y.-z., Liu, Z.-x., Li, Y.-s., Wang, X.-j., Lian, X.-r. and Zhou, Y.-q. (2023). Candidate gene localization of ZmDLE1 gene regulating plant height and ear height in maize.

